# Heterosynaptic interactions between dorsal and ventral hippocampus in individual medium spiny neurons of the nucleus accumbens ventromedial shell

**DOI:** 10.1101/2025.06.23.661109

**Authors:** Ashley E. Copenhaver, Sydnee Vance, Sarah A. Snider, Kaela Befano, J. Branwen She, Tara A. LeGates

## Abstract

Establishing learned associations between rewarding stimuli and the context under which those rewards are encountered is critical for survival. Hippocampal input to the nucleus accumbens (NAc) provides important environmental context to reward processing to support goal-directed behaviors. This connection consists of two independent pathways originating from the dorsal (dHipp) or ventral (vHipp) hippocampus, which have previously been considered functionally and anatomically distinct. Here, we show overlap in dHipp and vHipp terminal fields in the NAc, leading us to reconsider this view and raise new questions regarding the potential interactions between dHipp and vHipp pathways in the NAc. Using optogenetics, electrophysiology, and transsynaptic labeling in mice, we investigated anatomical and functional convergence of dHipp and vHipp inputs in the NAc. Transsynaptic labeling revealed a subpopulation of dually innervated cells in the NAc medial shell, confirmed by independent optogenetic manipulation of dHipp and vHipp inputs during whole-cell electrophysiological recordings. Further analysis revealed closely apposed dHipp and vHipp inputs along dendritic branches, and simultaneous stimulation of both inputs elicited heterosynaptic potentiation. Comparison of observed and theoretical success rates suggests interactions between dHipp-NAc and vHipp-NAc synapses may occur presynaptically. Altogether, these results demonstrate that inputs originating from dHipp and vHipp converge onto a subset of NAc neurons with synapses positioned to enable rapid heterosynaptic interactions, indicating integration of these inputs at the single-neuron level. Exploring the physiological and behavioral implications of this convergence will offer new insights into how individual neurons incorporate information from distinct inputs and how this integration may shape learning.

**SIGNIFICANCE STATEMENT:** Forming associations between rewards and the circumstances under which they are experienced is vital for survival. Hipp input to the NAc is essential for associating rewards with their environmental context to effectively guide motivated behaviors. This connection consists of two separate pathways originating from dHipp and vHipp that have long been considered distinct. Here, we reveal a subpopulation of neurons in the NAc shell innervated by both Hipp subregions as well as heterosynaptic interactions that occur between dHipp and vHipp synapses. These findings suggest that integration of distinct hippocampal information occurs at the single-neuron level, providing a critical mechanism underlying learning and motivated behavior while also opening new avenues for understanding how diverse contextual and reward signals shape decision-making.

## INTRODUCTION

The integration of diverse information across neural systems is crucial for driving complex behaviors. The brain dynamically integrates information from multiple sources, enabling organisms to react to stimuli and establish learned associations crucial for survival. Plasticity of and interactions between synapses enables coordinated regulation across multiple inputs, critical for orchestrating cell-wide functional responses (Bailey et al., 2000a; Govindarajan et al., 2006a; Harvey & Svoboda, 2007a; Redondo & Morris, 2011a; Schuman & Madison, 1994a). However, the mechanisms underlying how convergent inputs interact to shape neuronal responses remain unclear.

The nucleus accumbens (NAc) is a key brain region that integrates diverse information from several brain areas to effectively mediate goal-directed behaviors (Francis & Lobo, 2017; French & Totterdell, 2002; Gruber et al., 2009; Mannella et al., 2013; Marinescu & Labouesse, 2024; Salimpoor et al., 2013; Soares-Cunha et al., 2020). Hippocampal (Hipp) input is a major source of excitatory drive that influences NAc neuron activity (Brog et al., 1993; Calhoon & O’Donnell, 2013; Groenewegen et al., 1987; Kelley & Domesick, 1982; Li et al., 2018; O’Donnell & Grace, 1995; Phillipson & Griffiths, 1985; Trouche et al., 2019), and because of the Hipp’s essential role in contextual and spatial learning (Duarte-Guterman et al., 2015; Lamsa & Lau, 2019; Safari et al., 2021; Savage, 2004; Squire, 1992), Hipp-NAc synapses are also key sites for integrating rewards with related contextual and spatial information. Furthermore, the modulation of the strength of Hipp-NAc synapses is a critical mediator of motivated behaviors and is necessary for supporting contextual- and spatial-based reward learning (Huntley et al., 2020; Ibrahim et al., 2024; Ito et al., 2008, 2008; LeGates et al., 2018; Patterson et al., 2025, p. 202; Pennartz et al., 2011; Sjulson et al., 2018; Trouche et al., 2019; F.-C. Yang & Liang, 2014).

The connection between the Hipp and NAc consists of two independent pathways originating from the dorsal and ventral Hipp (dHipp and vHipp), which are considered functionally and anatomically distinct (Amaral & Witter, 1989; Bannerman et al., 2002; Fanselow & Dong, 2010; Friedman et al., 2002; K. B. Kjelstrup et al., 2008; K. G. Kjelstrup et al., 2002; Roberts et al., 2007; Strange et al., 2014). Notably, the NAc is one of the few regions that receives input from both the dHipp and vHipp (Friedman et al., 2002; Kelley & Domesick, 1982; Phillipson & Griffiths, 1985; Trouche et al., 2019). Although dHipp and vHipp are interconnected within the Hipp (Ishizuka et al., 1990; Swanson & Cowan, 1977; Tao et al., 2021), interactions through connections with other brain regions have been proposed as a mechanism linking cognitive and emotional processes (Fanselow & Dong, 2010). In a multifaceted environment, integration of diverse information is important for properly mediating complex behaviors, and the behavioral complexity governed by these pathways warrants synaptic integration to support learning and memory. This is supported by recent findings demonstrating a key role for convergence of dHipp and vHipp input to the amygdala in observational contextual fear memory (Terranova et al., 2022), yet the synaptic basis of this convergence and whether it is also observed in the NAc remains unknown.

Using a combination of microscopy, electrophysiology, and optogenetics, we identified and characterized a subpopulation of medium spiny neurons (MSNs) in the medial NAc shell (NAcSh) that are functionally innervated by both the dHipp and vHipp. dHipp and vHipp synapses on dually innervated MSNs have similar basal synaptic properties and are positioned at distances close enough to enable heterosynaptic interactions. These findings suggest that convergent Hipp inputs interact at the synaptic level, which may contribute to the integration of spatial and contextual information to modulate motivated behaviors. Overall, understanding the synaptic integration of dHipp and vHipp inputs in the NAc not only enhances our knowledge of reward-related behaviors, but also offers insight into the fundamental task of neuronal processing and provides a foundation for developing targeted interventions for disorders where these mechanisms are dysregulated.

## METHODS

### Animals

Male and female C57BL/6J, Drd1a-tdTomato, Ai14, and Ai195 mice were bred in-house. Mice were housed with same-sex cage mates in a temperature- and humidity-controlled environment under a 12:12 light cycle (lights on at 07:00), with food available *ad libitum*. We did not track estrous cycle in females. All experiments were performed in accordance with the regulations set forth by the Institutional Animal Care and Use Committee at the University of Maryland, Baltimore County.

### Stereotaxic surgeries

Mice (6-8 weeks old) were anesthetized with 3% isoflurane and underwent stereotaxic surgery. A Hamilton syringe was used to inject adeno-associated viruses (AAV) into the dHipp (in millimeters: from bregma, anterior-posterior (AP), -3.7; medial-lateral (ML), ±2.4; dorsal-ventral (DV), -2.8), vHipp (from bregma, AP, -3.7; ML, ±3.0; DV, -4.8), and NAc (in millimeters: from bregma, anterior-posterior (AP), +1.8; medial-lateral (ML), ±0.6; dorsal-ventral (DV), -4.7). Viruses were injected unilaterally or bilaterally and were infused at a rate of 0.1uL per minute. The injection needle was left in place for 10 minutes following the infusion. After surgery, mice were placed in their home cage on a heating pad for recovery. Carprofen (5mg/kg) was subcutaneously injected daily for 3 days after surgery. Mice recovered for 5-7 weeks for viral expression before anatomical or electrophysiological experiments. Viral constructs pAAV-hSyn-EGFP (Addgene #50465-AAV5) and pAAV-hSyn-mCherry (Addgene #114472-AAV5) were used to visualize dHipp and vHipp axon terminal fields. Anatomical labeling of dually innervated neurons involved the use of pENN.AAV.hSyn.Cre.WPRE.hGH (AAV1, 1013 titer, UMB Viral Vector Core), pAAV-EF1a-Flpo (AAV1, 1013 titer, UMB Viral Vector Core), and pAAV-hSyn-DIO-EGFP (Addgene #50457-AAV2). Specifically, pENN.AAV.hSyn.Cre.WPRE.hGH and pAAV-EF1a-Flpo were injected into the dHipp or vHipp of Ai195 mice for selective labeling of dually innervated neurons. pENN.AAV.hSyn.Cre.WPRE.hGH and pAAV-EF1a-Flpo were injected into the dHipp or vHipp and pAAV-hSyn-DIO-EGFP was injected in the NAc of Ai14 mice to label neurons single and dually innervated neurons. For optogenetic experiments, viral vectors pAAV-Syn-ChrimsonR-tdT (Addgene ID: 59171-AAV5) and either pAAV-Syn-Chronos-GFP (Addgene ID: 59170-AAV5) or pAAV-hSyn-hChR2(H134R)-EYFP (Addgene ID: 26973-AAV5) were used. For all experiments, injection sites were verified by fluorescence microscopy, and brains with off target expression were removed from the experiment.

### Visualization of axon terminal fields and dually innervated neurons

Mice expressing viruses for anatomical tracing experiments were first anesthetized with isoflurane and then perfused with 1X phosphate buffer solution (PBS), followed by 4% paraformaldehyde (PFA). Brains were removed and post-fixed in 4% PFA overnight at 4°C and were then transferred to 1X PBS and stored at 4°C for up to 3 weeks. Brains were sectioned (100μm) using a VT1000S Leica Microsystems vibratome. Slices were then mounted with Vectashield Antifade Mounting Medium on slides for imaging.

Images were captured with either a Zeiss LSM 900 Confocal microscope or Zeiss Axio Zoom.V16, both equipped with Zeiss software. To prevent cross talk among channels, images were obtained using sequential scanning to detect multiple fluorophores. To reduce background noise and graininess of images, 2x frame-averaging was used and gain was kept at or below 700. The laser powers used were: EGFP <15% and mCherry or tdTomato <15%. For imaging of dendritic segments, z-stacks with intervals of 0.2μm were acquired and further analyzed using Imaris.

To quantify individual and dually innervated neurons, images of dorsomedial NAcSh, ventromedial NAcSh, ventrolateral NAcSh, and ventromedial NAc core were captured on Zeiss LSM 900 Confocal microscope at 40x magnification, equipped with Zeiss software. Images were captured throughout the dorsomedial NAcSh, ventromedial NacSh, ventrolateral NacSh, and ventromedial NAc core. Cells were then manually counted and recounted a day apart. MSNs were identified based on soma size and the presence of highly branched, spiny dendrites, as compared to interneurons, which have larger somas and fewer spines. Analysis and counting were done by an experimenter blinded to conditions.

### Acute brain slice preparation for electrophysiology

Acute coronal slices containing the nucleus accumbens or hippocampus were prepared for whole-cell patch-clamp electrophysiology. Hipp-containing slices were kept to verify injection sites before proceeding with electrophysiological experiments in the NAc. Animals were deeply anesthetized with isoflurane, decapitated, and brains were quickly dissected and submerged in ice-cold, bubbled (carbogen: 95% O_2_/5% CO_2_) *N*-methyl-D-glucamine (NMDG) recovery solution containing the following (in mM): 93 NMDG, 2.5 KCl, 1.2 NaH_2_PO_4_, 11 glucose, 25 NaHCO_3_, 1.2 MgCl_2_, and 2.4 CaCl_2,_ pH=7.3-7.4, osmolarity=300-310 mOsm. Using a vibratome (VT1000S, Leica Microsystems), coronal slices (400 µm) were cut in cold, oxygenated NMDG. Slices were transferred to 32-34°C NMDG for 7-12 minutes to recover and were then transferred to room-temperature artificial cerebrospinal fluid (aCSF) containing the following (in mM): 120 NaCl, 3 KCl, 1.0 NaH_2_PO_4_, 20 glucose, 25 NaHCO_3_, 1.5 MgCl_2_·7H_2_O, and 2.5 CaCl_2,_ pH=7.3-7.4. Slices were allowed to recover for 1-hour at room-temperature before beginning electrophysiological recordings.

### Whole-cell recordings

We performed whole-cell patch-clamp recordings using an Axopatch 200B amplifier (Axon Instruments, Molecular Devices) and a Digidata 1550B digitizer (Axon Instruments). Slices were placed in a submersion-type recording chamber and superfused with room-temperature aCSF (flow rate 0.5-1mL/min). Patch pipettes (4-8MΩ) were made from borosilicate glass (World Precision Instruments) using a Sutter Instruments P-97 model puller. Cells were visualized using a 60x water immersion objective (Nikon Eclipse FN-1). All recordings were performed in voltage-clamp conditions from MSNs in the NAc ventromedial shell.

470nm and 635nm light was used for optogenetic stimulation of Hipp axon terminals in the NAc to record evoked excitatory postsynaptic currents (EPSC). We used light intensities ≤0.5mW to prevent non-specific activation of Chrimson by 470nm light as shown in Figure S1a-b, which aligns with other published studies (Christoffel et al., 2021; Klapoetke et al., 2014; Xia et al., 2020). Similar light intensity ranges were used for 635nm stimulation, and we did not observe non-specific stimulation with this wavelength at any light intensity (Figure S1c-d).

For experiments determining I-V relationship and AMPA:NMDA receptor ratios, the patch pipette solution was composed of 135 mM CsCl, 2 mM MgCl6-H2O, 10 mM HEPES, 4 mM Mg-ATP, 0.3 mM Na2-GTP, 10 mM Na2-phosphocreatine, 1 mM EGTA, 5 mM QX-314 bromide, and 100 μM spermine; pH=7.3-7.4; osmolarity=285-295mOsm. EPSCs were collected from holding potentials ranging from -80mV to +40mV to create an I-V curve. To calculate AMPA:NMDA receptor ratios, the AMPA component was defined by the peak amplitude at -70mV while the NMDA component was defined by the amplitude at +40mV at 50ms after stimulation. Paired pulse stimulation (100ms ISI) of dHipp or vHipp terminals was alternated every 10s to record EPSCs at dHipp-MSN and vHipp-MSN synapses. PPR was determined by calculating 5-minute averages of EPSC amplitudes from our paired pulses (EPSC1 and EPSC2) and dividing EPSC2 by EPSC1 (ie: EPSC2/EPSC1 = PPR). For all other experiments, patch pipettes were filled with a solution containing 130 mM K-gluconate, 5 mM KCl, 2 mM MgCl6-H2O, 10 mM HEPES, 4 mM Mg-ATP, 0.3 mM Na2-GTP, 10 mM Na2-phosphocreatine, and 1 mM EGTA; pH=7.3-7.4; osmolarity=285-295mOsm. We used the following exclusion criteria to eliminate unhealthy cells and unreliable recordings: 1) We only proceeded with experiments on cells with series resistances <10MΩ, 2) Cells were excluded if their series resistance changed by >20% (comparing the resistance at the beginning and end of the experiment), 3) Cells in poor health or poor recording status were excluded (i.e. cell partially or fully sealed up, a decrease in holding current >100pA that is consistent with the cell dying, an increase in jitter post HFS, and/or an increase in response failure rate to > 50%).

### Visualization of dHipp/vHipp terminals with dually innervated neurons

Using a small subset of slices from electrophysiological experiments, dually innervated MSNs were patched and filled with CF633 or CF647 dye (Biotium). Upon completion of electrophysiology experiments, the 400μm-thick slices were fixed in 4% PFA for 4-8 hours at 4°C. Slices were then mounted in Invitrogen ProLong Glass Antifade Mountant (Thermo Fisher Scientific, #P36980) on slides with spacers (400μm-thick) for imaging. Images were captured with a Zeiss LSM 910 confocal microscope equipped with Zeiss software. To prevent cross talk among channels, images were obtained using sequential scanning to detect multiple fluorophores. To reduce background noise and graininess of images, 2x frame-averaging was used and gain was kept at or below 700. The laser powers used were: EGFP <15%, mCherry or tdTomato <15%, CF633 <10%. For imaging of dendritic segments and analysis of synapse distributions, z-stacks (0.2μm slices) of distal dendrites were acquired. Surface rendering was conducted using Imaris to create 3-dimensional reconstructions of dHipp and vHipp inputs and MSN dendrites. These images were then analyzed in Imaris to identify putative dHipp-MSN and vHipp-MSN synapses and calculate Eucledian distances (edge-to-edge) between nearest neighboring dHipp-MSN and vHipp-MSN synapses along 100μm dendritic segments.

### Quantification, statistical analysis, and reproducibility

We found no statistically significant difference between males and females, so data from male and females are plotted together. Both the number of cells and number of mice are reported for each experiment. Plots represent the mean ± SEM. The sample size (n) per condition represents the number of cells unless otherwise indicated in the figure caption. Potentiation scores were calculated as the amplitude of the EPSC evoked from synchronous stimulation divided by the sum of the amplitudes from the alternating stimulations. All statistical analyses were performed using Graphpad Prism 9/10 software. Statistical significance (p<0.05) was assessed using parametric or non-parametric versions of paired t-tests, or one-way or two-way ANOVA with multiple comparisons. Distributions of nearest neighbor distances were compared to a uniform distribution using a Chi-square goodness of fit test. For box plots, the line in the middle of the box is plotted at the median. The box extends from the 25th to 75th percentiles. Whiskers represent minimum and maximum, and individual data points are shown. Figure schematics and experimental timelines were created with BioRender.com.

## RESULTS

### dHipp and vHipp axon terminal fields overlap in the NAc medial shell

Using viral-mediated labeling to visualize Hipp neurons and their terminal fields, we examined innervation of the NAc by the posterior dHipp and vHipp. We injected an adenoassociated virus (AAV) expressing eYFP into the dHipp and an AAV expressing mCherry into the vHipp (Figure 1a-b). After allowing 5-7 weeks for expression, brains were fixed, sectioned, and imaged using confocal microscopy. We observed dHipp axon terminal fields in the NAc core and lateral shell while the vHipp axon terminal fields were primarily found in the medial shell, in line with previous work (Figure 1c). However, we also observed strong overlap of dHipp and vHipp axon terminal fields in the medial NAcSh (Figure 1c-d), which aligns with the results presented by Brog et al., 1993. 3-D reconstruction of z-stack images of these overlapping axon fields revealed adjacent dHipp and vHipp axons (Figure 1d), highlighting the close proximity of dHipp and vHipp afferents within the medial NAcSh.

**Figure 1.**
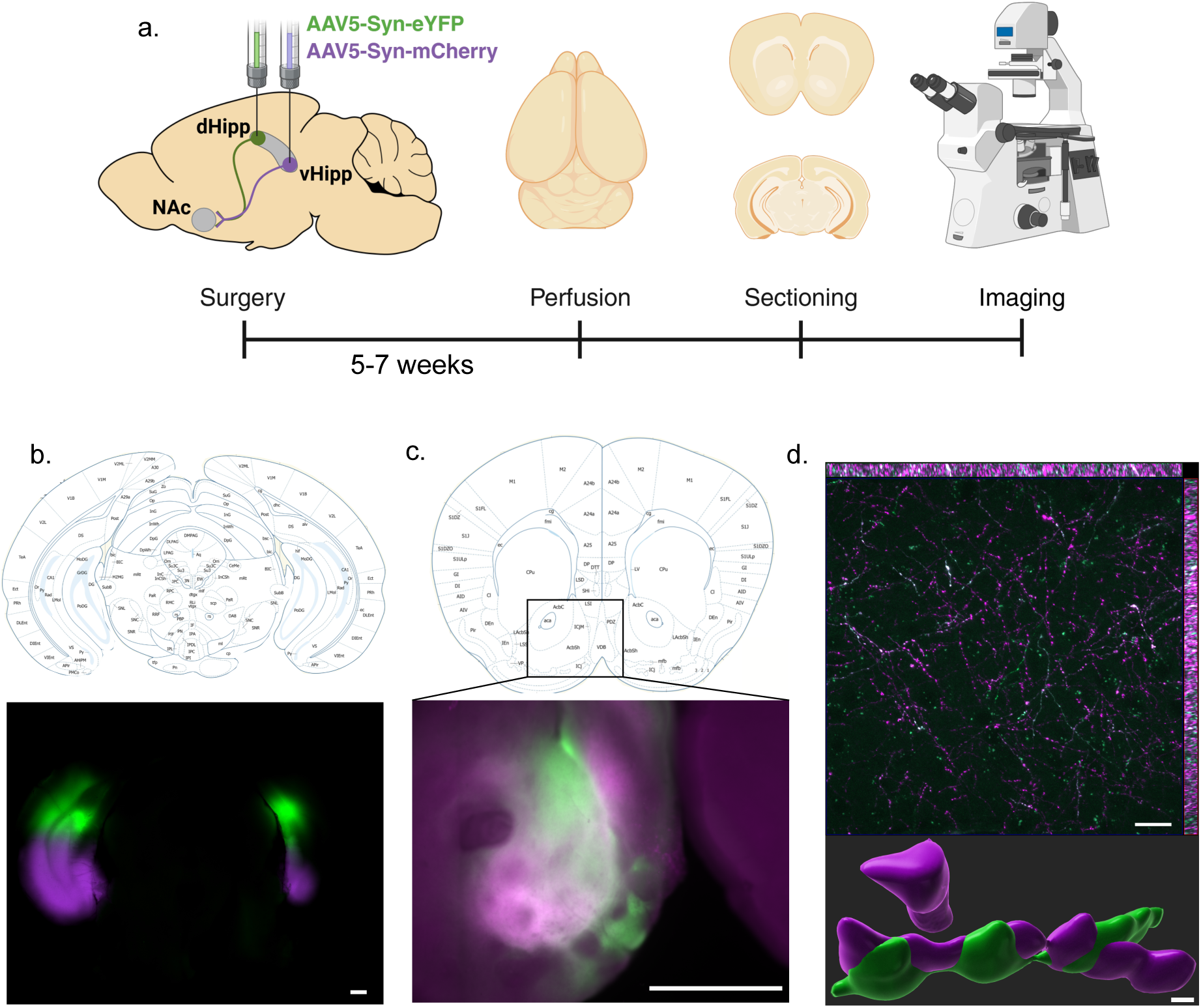
Overlapping dHipp and vHipp axon terminal fields in the NAc. a) Viral injection strategy and experimental timeline. b) Injection sites in the hippocampus. Scale bar = 500µm. Atlas image from (Paxinos & Franklin, 2019) c) Overlap of dHipp and vHipp axon terminal fields in the NAc. Scale bar = 500µm. Atlas image from (Paxinos & Franklin, 2019) d) Close-up image of overlapping dHipp and vHipp axon terminal fields within the NAc ventromedial shell (top). Scale bar = 20µm. With 3-dimensional reconstruction of overlapping axon terminals (bottom). Scale bar = 0.5µm

### Identification of NAc neurons dually innervated by the dHipp and vHipp

Given the apposition of dHipp and vHipp terminals, we sought to determine whether individual neurons were innervated by both hippocampal regions. To do this, we utilized transsynaptic expression of Cre and Flp recombinases to selectively target NAc neurons innervated by dHipp and vHipp pathways (Figure 2a). We injected AAV1-hSyn-Cre into the dHipp and AAV1-Ef1a-Flpo into the vHipp of Ai195 mice in which GcaMP6s is only expressed in the presence of Cre and Flp. This approach allowed for targeted expression of GCaMP6s only in neurons that are innervated by both dHipp and vHipp (Figure 2b). After allowing 5-7 weeks for expression, brains were fixed, sectioned and imaged with confocal microscopy using GCaMP6s to visualize dually innervated neurons. Similar to our observations of terminal field overlap, we observed GCaMP6s-expressing cells in the medial NAcSh (Figure 2c), suggesting dual innervation of NAc neurons by both dHipp and vHipp. Furthermore, examination of GCaMP6s-expressing cells revealed morphology similar to MSNs (based on soma size and spiny, highly branched dendrites) (Figure 2d), suggesting that individual NAc MSNs receive input from both dHipp and vHipp.

**Figure 2.**
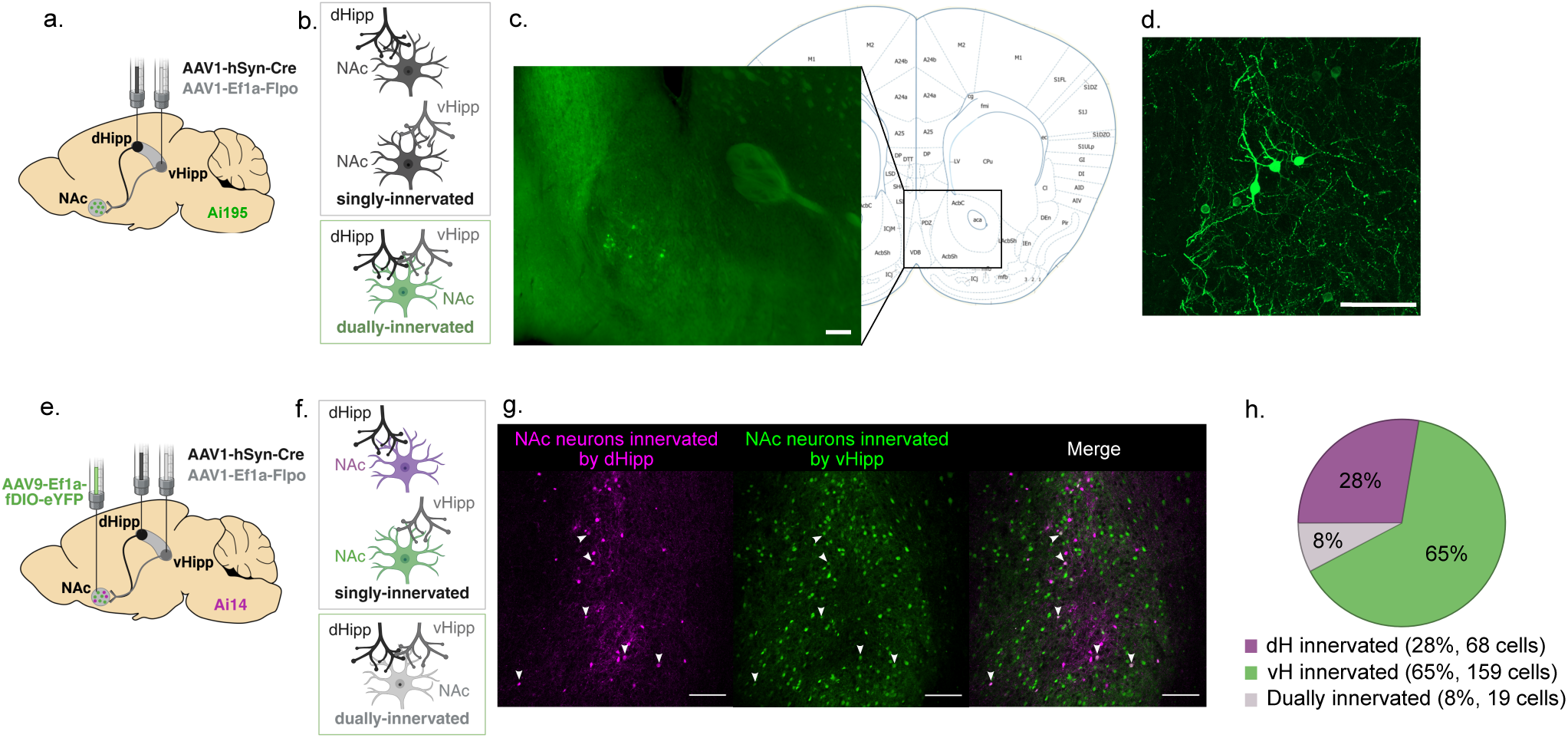
dHipp^+^/vHipp^+^-innervated neurons in the medial NAc shell. a) Viral targeting strategy to selectively label dually innervated neurons b) Cartoon representation of singly vs dually innervated neurons. c) Image of medial NAcSh depicting GCaMP-expressing cells. Scale bar = 100µm. Atlas image from (Paxinos & Franklin, 2019) d) Close-up image of GCaMP-expressing cells in the medial NAcSh, where neuron morphology is consistent with medium spiny neurons. Scale bar = 100µm. e) Viral targeting strategy and experimental timeline to label cells innervated by dHipp and vHipp. f) Cartoon representation of singly vs dually innervated neurons. g) Image of cells innervated by dHipp (green) and vHipp (magenta). Dually innervated neurons express both reporters and appear white. Arrowheads point out examples of dually innervated neurons. Scale bar = 100µm h) Quantification of cells innervated single and dually innervated neurons in the medial NAcSh across 4 slices.

To gain better insight into the proportions of cells innervated by dHipp, vHipp, or both, we used our transsynaptic strategy to differentially label cells innervated by dHipp and vHipp. Specifically, we injected AAV1-hSyn-Cre into the dHipp and AAV1-Ef1a-Flpo into the vHipp of Ai14 mice in which there is Cre-dependent expression of tdTomato. We also injected AAV9-Ef1a-fDIO-eYFP into the NAc of these mice (Figure 2e). As a result, cells innervated by dHipp express tdTomato, cells innervated by vHipp express eYFP, and those innervated by both will express both reporters (Figure 2f-g). Quantification of cells in medial NAcSh revealed approximately 8% of cells were innervated by dHipp and vHipp (dually innervated). The remainder of cells were innervated by one Hipp subregion (singly innervated). Of these, 28% cells were innervated by dHipp, 65% were innervated by vHipp (Figure 2h). This suggests that a small proportion of neurons in the medial NAcSh are dually innervated. However, it is worth noting that previous work has reported low efficiency of transsynaptic transport, leading us to believe that our visualization strategy underrepresents the number of dually innervated neurons (Zingg et al., 2017).

To independently verify whether individual NAc neurons were dually innervated and determine whether our anatomical results translated into functional convergence, we used optogenetics and whole-cell electrophysiology to independently stimulate dHipp and vHipp inputs while recording from individual NAc neurons. We focused on MSNs because they comprise 95% of the NAc neuron population, align with the morphology of cells identified by our anatomical tracing studies (Figure 2), and have been shown to receive functional input from either dHipp or vHipp in separate studies. To independently stimulate dHipp and vHipp inputs, we used a dual wavelength optogenetic strategy involving viral expression of Chronos and Chrimson, which are maximally sensitive to different wavelengths of light (480nm and 660nm, respectively) (Klapoetke et al., 2014). AAV2-Syn-Chronos-eYFP or AAV2-Syn-Chrimson-mCherry were injected into the dHipp or vHipp. At least 5 weeks after surgery, coronal brain slices containing the NAc were prepared for electrophysiology. Using whole-cell patch-clamp electrophysiology, we recorded from MSNs in the medial NAcSh and used alternating 473 and 635nm light to measure dHipp and vHipp evoked responses in the MSNs. Keeping the light intensity below 1mW prevented cross-contamination, allowing us to measure and independently manipulate dHipp and vHipp inputs with 473 and 635nm light (Figure S1 and 3a). Using this strategy, we identified a subpopulation of MSNs in the medial NAcSh that responded to both dHipp and vHipp stimulation, demonstrating dual functional innervation of individual NAc MSNs by both dHipp and vHipp (Figure 3a). Dually innervated neurons accounted for ∼20-50% of neurons surveyed, a greater proportion than that observed with our transsynaptic labeling (Figure 2). The variation in dually innervated neurons appeared to depend on recording site, with higher proportions detected in regions that overlapped with those identified by our labeling approach.

**Figure 3.**
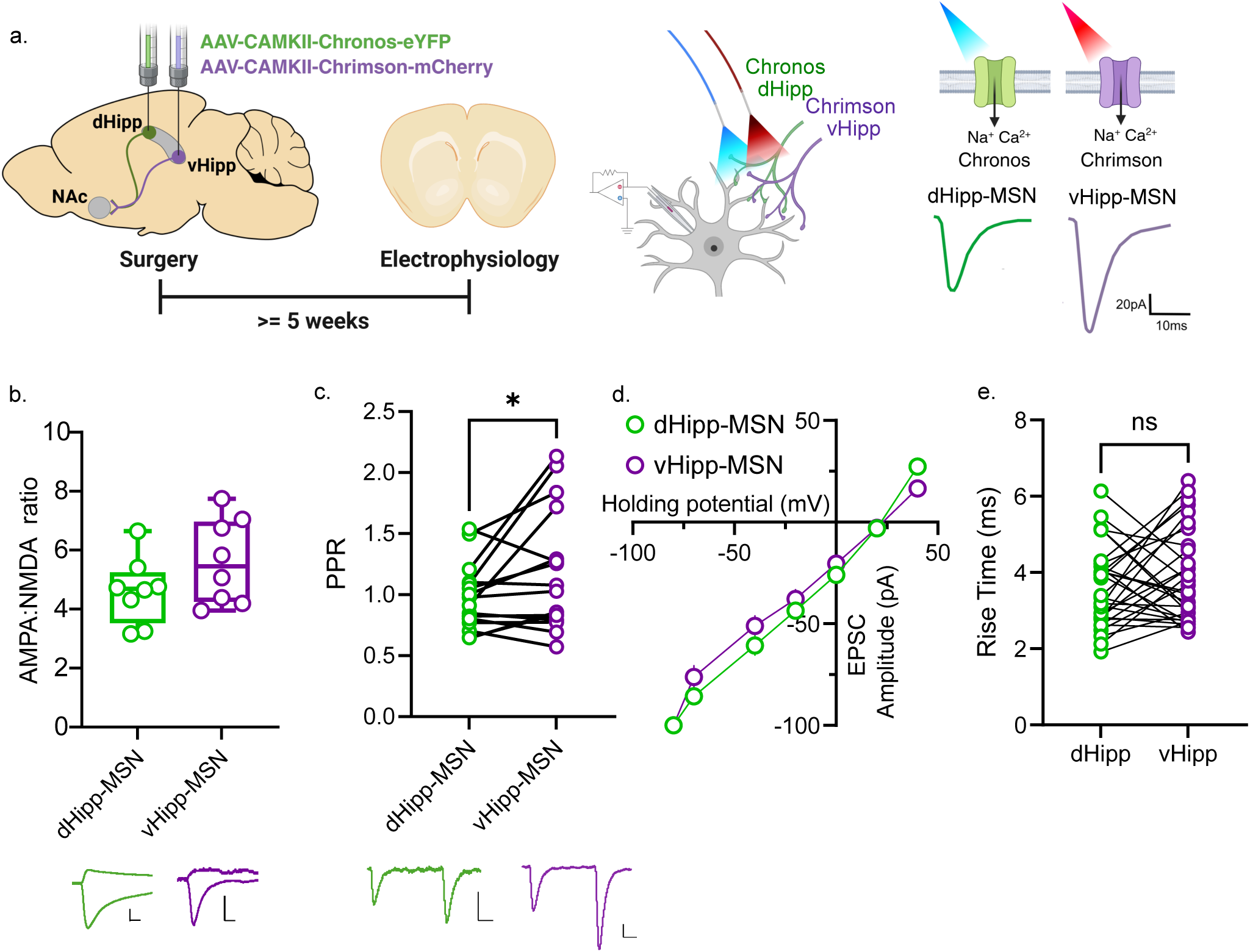
Individual NAc neurons are functionally innervated by dHipp and vHipp. a) Viral targeting and optogenetic approach. Electrophysiological recording strategy and representative traces from a dually innervated NAc MSN. b) Similar AMPA:NMDA receptor ratios between dHipp-and vHipp-MSN synapses (n=8 cells/4 mice, *p*=0.1953, Wilcoxon test). Scale bars on representative traces = 40pA/10ms. c) PPRs recorded from dHipp-MSN and vHipp-MSN synapses on dually innervated neurons (n=16 cells/10 mice, *p=0.0394, Paired t-test). Scale bars on representative traces = 20pA/20ms. d) Rectification curve reveals linear relationship, indicating a lack of calcium-permeable AMPA receptors (n=8 cells/4 mice). e) Rise time was calculated for dHipp- and vHipp-evoked EPSCs, showcasing similar rise times (n=31 cells/12 mice, *p*=0.2877, Wilcoxon test).

Synaptic properties and strengths influence how a neuron responds to and integrates information from distinct inputs, leading us to examine and compare the synaptic profiles of dHipp-MSN and vHipp-MSN synapses in dually innervated neurons. To examine excitatory synaptic strength, we measured AMPAR:NMDAR ratios at dHipp-MSN and vHipp-MSN synapses and found no difference, suggesting that the strength of dHipp- and vHipp-MSN synapses is similar (Figure 3b). We then used paired pulse stimulation, alternating paired pulses such that each pathway was stimulated every 20s, to examine paired-pulse ratios (PPR). PPR was significantly greater at vHipp-MSN synapses as compared to dHipp-MSN synapses (Figure 3c), suggesting potential differences in release probability, though this may be driven by a subpopulation of recorded neurons.

Calcium-permeable AMPA receptors (CP-AMPAR) have been identified at ventral subiculum-NAc synapses (Boxer et al., 2023) and have been implicated in NAc plasticity and related behaviors (Carr, 2020; Mameli et al., 2009; McCutcheon et al., 2011; Wolf & Tseng, 2012). Given their role in plasticity and behavior, we examined whether CP-AMPARs were present at dHipp-or vHipp-MSN synapses. Using their characteristic inward rectification and non-linear current-voltage relationship (Cull-Candy et al., 2006; Liu & Zukin, 2007), we performed electrophysiological experiments to identify the predominant population of AMPARs at dHipp and vHipp-MSN synapses. Our results revealed that neither Hipp-MSN synapse contained a significant population of CP-AMPARs (Figure 3d). Collectively, these results indicate that dHipp and vHipp synapses in dually innervated NAc MSNs have comparable basal postsynaptic properties with presynaptic differences in release probability.

We then evaluated the potential for synaptic or dendritic-level interactions between dHipp and vHipp synapses by comparing rise times and the spatial locations of putative dHipp and vHipp synapses. Our rise time analysis revealed similar rise times of dHipp- and vHipp-evoked responses, suggesting that dHipp-MSN and vHipp-MSN synapses are located a similar distance away from the cell body (Figure 3e). We filled a subset of cells electrophysiologically confirmed to be dually innervated with a fluorescent dye (Figure 4a), and slices were fixed and prepared for confocal microscopy. We used confocal microscopy and image analysis to measure the spatial location of potential dHipp and vHipp synapses along the dendrites of a dually innervated MSN using the eYFP co-expressed with Chronos, the mCherry co-expressed with Chrimson, and the dye. From this analysis, we found several examples of dHipp and vHipp puncta that are spatially close (<2µm) along distal dendrites of dually innervated MSNs (Figure 4b-c). We analyzed 3-dimensional reconstructions of z-stack images to identify putative dHipp and vHipp inputs on MSNs dendrites and conducted nearest neighbor analysis to measure the spatial relationships between dHipp-MSN and vHipp-MSN synapses. We found that many (>50%) dHipp-MSN and vHipp-MSN synapses are located within a 0-2µm distance (Figure 4d-e), suggesting that dHipp–MSN and vHipp–MSN synapses cluster preferentially near each other. Altogether, these results demonstrate functional innervation of individual MSNs by both dHipp and vHipp, where synapses are frequently observed in close spatial proximity along MSN dendrites.

**Figure 4.**
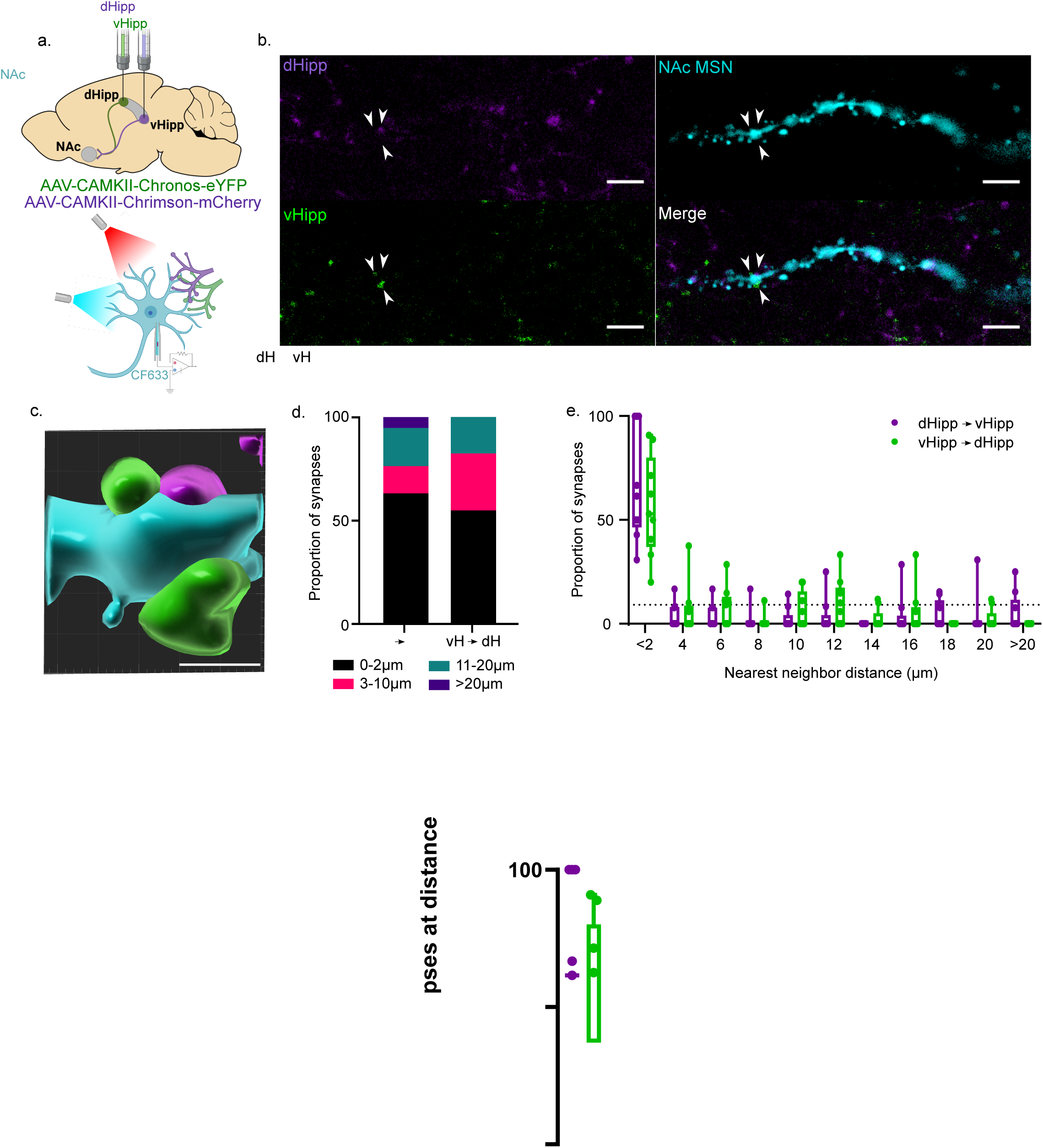
Spatial proximity of dHipp and vHipp puncta along distal dendrites of MSN. a) Cartoon depicting optogenetic, electrophysiology, and cell-filling strategy. b) Combined and separated images depicting an MSN filled with CF633 confirmed to have functional dHipp and vHipp synapses, dHipp puncta (green), vHipp puncta (red). Scale bar = 5µm. c) Three-dimensional reconstruction of an MSN dendritic segment (cyan) with overlapping/merged dHipp (red) and vHipp (green) afferents. Scale bar = 1µm. d) Summary data showing the proportions of nearest neighbor distances in a stacked bar graph. e) Nearest neighbor distances shown in 2µm bins (n=9 cells/6 mice). The observed distribution differed significantly from a uniform distribution (p<0.0001; Chi-square goodness-of-fit test). Dotted line indicates expected values from uniform distribution.

### Heterosynaptic interactions between dHipp and vHipp synapses in NAc MSNs

Synapses that are sufficiently close (<2µm apart) can interact and influence each other’s activity (Fu et al., 2012; Harvey & Svoboda, 2007a; Murakoshi et al., 2011; Xia et al., 2020; Yuste, 2011). Though activity in dHipp and vHipp is not typically temporally coordinated, learning has been found to increase synchrony between dHipp and vHipp (Biane et al., 2023; Morici et al., 2025). Synchronous activity has been shown to induce plasticity through heterosynaptic interactions between adjacent synapses, raising the question of whether the spatial proximity of dHipp and vHipp synapses facilitates heterosynaptic interactions. To address this, we alternated 473nm and 635nm stimulation for 2 minutes to measure EPSCs evoked from independent stimulation of each Hipp pathway. We then synchronized 473nm and 635nm stimulations for 2 minutes to record an evoked EPSC from the simultaneous stimulation of both dHipp and vHipp inputs (Figure 5a,b). If no heterosynaptic interactions occur, then the EPSC amplitudes from the alternating stimulation should be the linear sum of the EPSC amplitude evoked by simultaneous stimulation. Instead, our data show a supralinear summation of separately measured EPSCs (Figure 5b), indicating that heterosynaptic potentiation occurs between dHipp and vHipp synapses, an effect observed across multiple light intensities in most cells (Figure S2). We conducted the same experiment in cells that were not dually innervated and found no difference between EPSC amplitudes evoked by alternating and synchronous stimulation, demonstrating that this effect is specific to dually innervated neurons and not a nonspecific artifact resulting from cross-contamination (Figure 5c).

**Figure 5.**
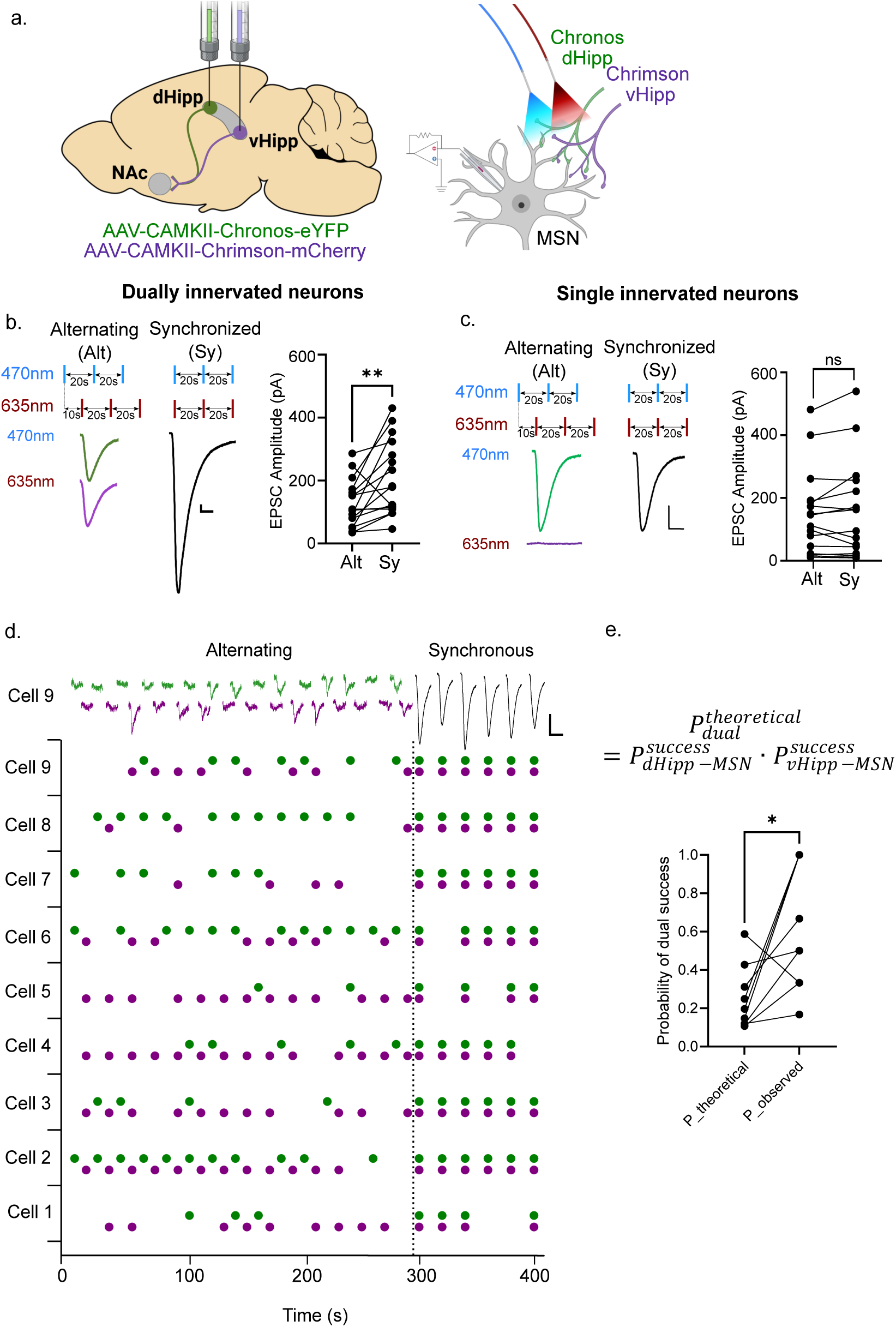
Heterosynaptic potentiation of dHipp-MSN and vHipp-MSN synapses. a) Viral expression, optogenetic, and electrophysiology strategy. b) Recording from dually innervated MSNs, sequential and synchronous stimulation assay reveals non-linear summation of responses (n=16 cells/10 mice, \*\**p*=0.0027, Wilcoxon test). Scale bars on representative traces = 20pA/10ms. c) Sequential and synchronous stimulation assay performed in singly innervated MSNs reveals similar responses to a single wavelength of light or both wavelengths of light (n=16 cells/10 mice, *p*=0.4037, Wilcoxon test). Scale bars on representative traces = 20pA/10ms. d) Raster plot showing failures and successes from minimal stimulation assay. Time period 300-420s is synchronized light pulses. Representative traces from Cell 9 are shown above the plot. Scale bars on representative traces = 30pA/20ms. e) Theoretical probability of dual success, *P_exp_*, compared with observed dual response success, *P_obs_*, during sync period (*n*=9 cells/5 mice, \**p*=0.0273, Wilcoxon test).

To better understand the site of synaptic interaction, we compared the failure rates in a separate experiment using minimal stimulation to evoke pathway-specific responses ∼50% of the time so that we could appropriately assess an increase or decrease in failure rate during synchronous stimulation (Figure 5d). We measured EPSCs evoked from alternating and synchronous stimulation of dHipp-MSN and vHipp-MSN synapses. We then calculated the theoretical probability of eliciting a response in both the dHipp- and vHipp-MSN pathways, *P_dual_*, if they were stimulated at the same time. Assuming the pathways are independent/non-interacting, we can calculate this probability as the product of their individual success properties: 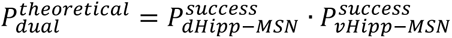. We compared this theoretical probability to the observed success rate of sync responses that were classified as dual-evoked responses, finding a significant difference in the observed and theoretical values (Figure 5e) suggesting presynaptic mechanisms may facilitate heterosynaptic interactions between dHipp- and vHipp-MSN synapses.

## DISCUSSION

The experiments in this study identified a subpopulation of neurons within the NAc that are dually innervated by dHipp and vHipp. These neurons are primarily located in the NAc ventromedial shell with morphological characteristics that align with those of MSNs. We found that dHipp-MSN and vHipp-MSN synapses in dually innervated neurons have similar basal synaptic properties including comparable release probabilities, synaptic strength, and expression of calcium-impermeable AMPARs. Examination of EPSCs evoked by individual or synchronous stimulation of dHipp and vHipp revealed supralinear summation of EPSCs, suggesting that the synchronous activation of dHipp and vHipp inputs induces short-term heterosynaptic potentiation of dHipp-MSN and vHipp-MSN synapses likely due to presynaptic mechanisms. Altogether, these results reveal mechanisms of dHipp and vHipp synaptic integration within the NAc medial shell, which may be key to supporting the integration of cognitive and emotional processes needed to effectively mediate goal-directed behaviors.

### Innervation of the NAc by dHipp and vHipp

Our viral-mediated axon terminal field labeling strategy revealed overlapping dHipp and vHipp innervation patterns within the NAc ventromedial shell. At first glance, this finding seems to contradict previous reports that have suggested that the dHipp and vHipp preferentially innervate distinct parts of the NAc. However, these reported innervation patterns appear to depend upon the anterior/posterior coordinate used for Hipp viral injections. Most studies have targeted the posterior vHipp and anterior dHipp, showing vHipp projections in the NAc core and medial shell, with dHipp projections in the NAc core and lateral shell (Groenewegen et al., 1987; Ibrahim et al., 2024; Li et al., 2018; Mulder et al., 1998; Scofield et al., 2016). But when the posterior dHipp was targeted with an anterograde tracer, dHipp axon terminal field expression was observed in the NAc ventromedial shell (Ibrahim et al., 2024). Furthermore, the injection of a retrograde tracer into the NAc ventromedial shell showed that neurons located in the posterior dHipp and vHipp were projecting to this area of the NAc (Groenewegen et al., 1987). Our findings align with the latter two studies, as we injected into the posterior Hipp (A/P: -3.7mm) and found overlapping dHipp and vHipp axon terminal fields and dually innervated neurons primarily in the NAc ventromedial shell.

Using a transsynaptic viral-mediated labeling technique, we were able to visualize a small subpopulation of dually innervated neurons that had morphological characteristics similar to MSNs (Figure 2). We confirmed this electrophysiologically, demonstrating that dHipp and vHipp inputs form synaptic connections with individual MSNs within the medial NAcSh (Figure 3). Because this labeling method has low transsynaptic efficiency (Zingg et al., 2017) and requires successful transfer of both Cre and Flp payloads, it likely underrepresents the number of dually innervated neurons. This is supported by our electrophysiology recordings, as within a single slice, we were able to record from a larger number of dually innervated neurons than were indicated from the anatomical tracing study. Further optimization of this transsynaptic approach could improve detection and determine whether sparsely distributed NAc interneurons also receive dual hippocampal input.

### dHipp and vHipp synapses have similar properties in dually innervated NAc neurons

We found that dHipp- and vHipp-MSN synapses contain primarily calcium-impermeable AMPARs and have similar excitatory synaptic strengths, as measured by AMPA:NMDA receptor ratios (Figure 3). These similarities suggest that, in basal conditions, both pathways may contribute equally to downstream circuit activity. Since the dHipp and vHipp convey different types of behavioral information, this may mean that Hipp-related contextual and sensory cues are equally capable of influencing motivated behaviors. While these similarities are present at baseline, several studies have highlighted changes that occur to dHipp-MSN and vHipp-MSN synapses in response to hormones, neuromodulators, cocaine, external stress, and experience (Floresco et al., 2001; LeGates et al., 2018; Peleg-Raibstein & Feldon, 2006; R. Yang & J. Mogenson, 1984; Sjulson et al., 2018; Williams et al., 2020).

### Synaptic interactions between distinct Hipp inputs

The NAc is a critical integrator of several major excitatory inputs, many of which exhibit coordinated or synchronous activation (Mahon et al., 2006; Stern et al., 1998). While dHipp and vHipp activity is not typically temporally coordinated, increased synchrony has been associated with learning (Biane et al., 2023; Morici et al., 2025). We found that dHipp-MSN and vHipp-MSN synaptic interactions during simultaneous stimulation elicit heterosynaptic potentiation (Figure 5). Convergent afferent activity and heterosynaptic interactions can alter plasticity thresholds or propagate plastic changes across synapses, fundamentally influencing neuronal information processing and storage (Chistiakova et al., 2015; Reissner et al., 2010). Several mechanisms underlying heterosynaptic plasticity have been identified, including neuromodulator-dependent forms, synaptic tagging and capture, and spatially clustered plasticity, each engaging distinct molecular pathways that ultimately promote plasticity across synapses (Bailey et al., 2000b; Govindarajan et al., 2006b; Harvey & Svoboda, 2007b; Redondo & Morris, 2011b; Schuman & Madison, 1994b). Within the NAc, convergent inputs interact at the synaptic level via a combination of presynaptic and postsynaptic mechanisms to influence behavioral output (O’Donnell & Grace, 1995; Xia et al., 2020; F.-C. Yang & Liang, 2014; Yu et al., 2022). Our analysis of observed and theoretical success rates suggests that dHipp/vHipp heterosynaptic potentiation may occur presynaptically (Figure 5). Axo-axonic synapses in the striatum are rare, but recent work has highlighted heterosynaptic interactions between the basolateral amygdala (BLA), medial prefrontal cortex (mPFC) and paraventricular nucleus of the thalamus (PVT) during co-activation that occur due to presynaptic innervation of mPFC and PVT by BLA (Kornhuber & Kornhuber, 1983; Xia et al., 2020). Additionally, co-activation of vHipp-MSN and BLA-MSN synapses induces long-term potentiation that requires the activity of dopaminergic terminals and D1 receptors (Yu et al., 2022). While this may be due to activity of postsynaptic receptors given that this form of plasticity is more prominent on D1-expressing MSNs, D1 receptors are also expressed presynaptically on glutamatergic terminals and astrocytes where their signaling can alter release probability (Corkrum et al., 2020; Nicola & Malenka, 1997; Zhang et al., 2014).

### Implications of integration that occurs at the synaptic level

The connection between the Hipp and NAc is crucial for integrating reward with related contextual and spatial information (Britt et al., 2012; Floresco & Phillips, 1999; Humphries & Prescott, 2010; Ito et al., 2008; Lansink et al., 2008, 2009; LeGates et al., 2018; Pennartz et al., 2011; Sjulson et al., 2018). Functional imaging studies in humans have revealed greater Hipp-NAc connectivity associated with positive reinforcement learning (Davidow et al., 2016), increased reward seeking in adolescents (Huntley et al., 2020), and greater positive affect in adults (Heller et al., 2020), implicating a strong link between the strength of Hipp-NAc connectivity and reward-related behaviors. Similar findings have been demonstrated in rodent models which examined Hipp-NAc connectivity using *in vivo* electrophysiology during spatial and contextual-based reward tasks (Sjulson et al., 2018; Sosa et al., 2020; Trouche et al., 2019). The causal relationship between Hipp-NAc connectivity and reward behaviors has been established with lesion and optogenetic studies, demonstrating that Hipp-NAc synapses are required for contextual- and spatial-based reward tasks (Floresco & Phillips, 1999; Ito et al., 2008, p. 200; LeGates et al., 2018; Trouche et al., 2019; Zhou et al., 2019). This includes our previously published work, which demonstrated that the strength of vHipp-NAc synapses is a critical mediator of reward-related behaviors (LeGates et al., 2018).

dHipp and vHipp convergence within other areas of the brain has been proposed to be a key circuit mechanism connecting cognitive and emotional processes (Fanselow & Dong, 2010) and may be crucial to establishing and retrieving memories. The dHipp and vHipp are known to convey distinct information critical for driving NAc-mediated motivated behaviors (Ibrahim et al., 2024; Ito et al., 2008; LeGates et al., 2018; Muir et al., 2020; Patterson et al., 2025; Sjulson et al., 2018; Sosa et al., 2020; Trouche et al., 2019; Williams et al., 2020; A. K. Yang et al., 2020; F.-C. Yang & Liang, 2014). Therefore, the convergence and plasticity we observed may provide a basis by which distinct information conveyed by dHipp and vHipp can shapes the NAc’s response to more nuanced contextual and associative information. Similar convergence has been observed in the amygdala, where dHipp and vHipp activate and reactivate, respectively, specific neuron ensembles to mediate contextual fear memory (Terranova et al., 2022). Since Hipp input exerts a strong influence over NAc neuron activity and information flow, it is likely these interactions encourage integration of distinct information conveyed by the dHipp and vHipp, critically impacting behavioral outputs.

Altogether, the results presented here provide evidence for integration of dHipp and vHipp information within individual MSNs of the NAc ventromedial shell. The combination of techniques enabling identification of similar synaptic profiles and rapid interactions occurring between dHipp-and vHipp-MSN synapses provides a foundation for similar studies investigating the integration of distinct inputs at the level of individual neurons. Ultimately, these findings advance our understanding of how spatial and contextual information may contribute to reward-related behaviors while also raising further questions regarding how each of these distinct pathways are regulated.

## Acknowledgments

We would like to thank Dr. Tagide deCarvalho for her support with microscopy and image analysis, and Dr. Ramesh Chandra for his help with viruses. We would also like to thank Jennifer Pham for her help with injections and Dr. Brian Mathur for his helpful suggestions on this project. This work was supported by NSF-IOS2402645, T32GM144876-02, T34GM136497, the Merck Academic Fellowship Program, and startup funds provided by UMBC.

---

**Figure S1.**
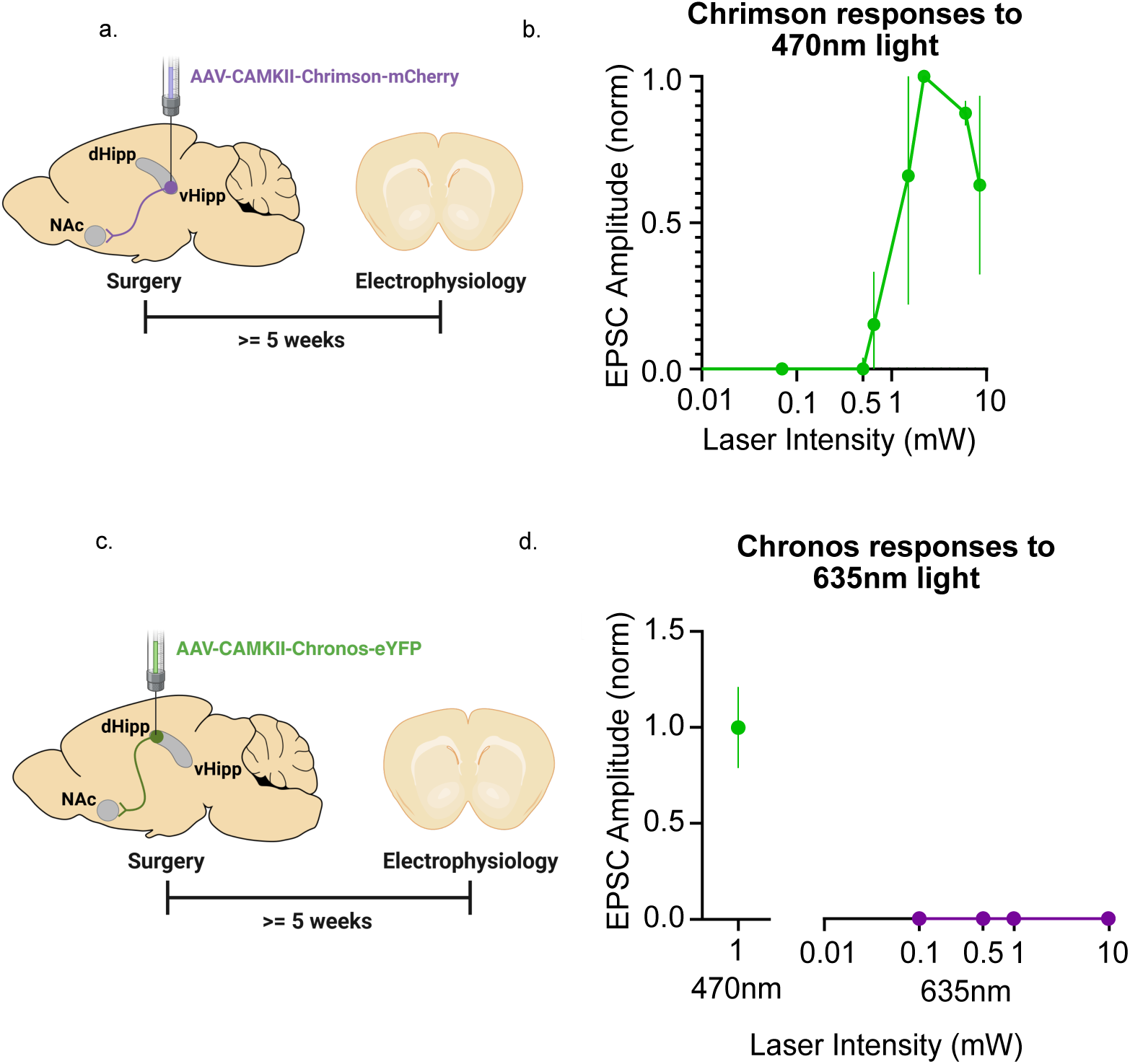
Prevention of cross-contamination when using 470nm light and the optogenetic protein, Chrimson. a) Mice were injected with a Chrimson-containing viral construct and slices were taken at least 5 weeks after surgery to perform electrophysiology. b) A response curve was generated by sampling Chrimson-evoked responses using 470nm light, showing minimal responses at a laser intensity of 0.75mW, and no Chrimson-evoked response at laser intensities ≤0.5mW. EPSC amplitudes were normalized to the maximum EPSC amplitude. c) Mice were injected with a Chronos-containing viral construct and slices were taken at least 5 weeks after surgery to perform electrophysiology. d) No responses were observed at any of the sampled light intensities. Responses were also recorded in response to 470nm (1mW) to verify successful targeting and stimulation (shown in green on the left).

**Figure S2.**
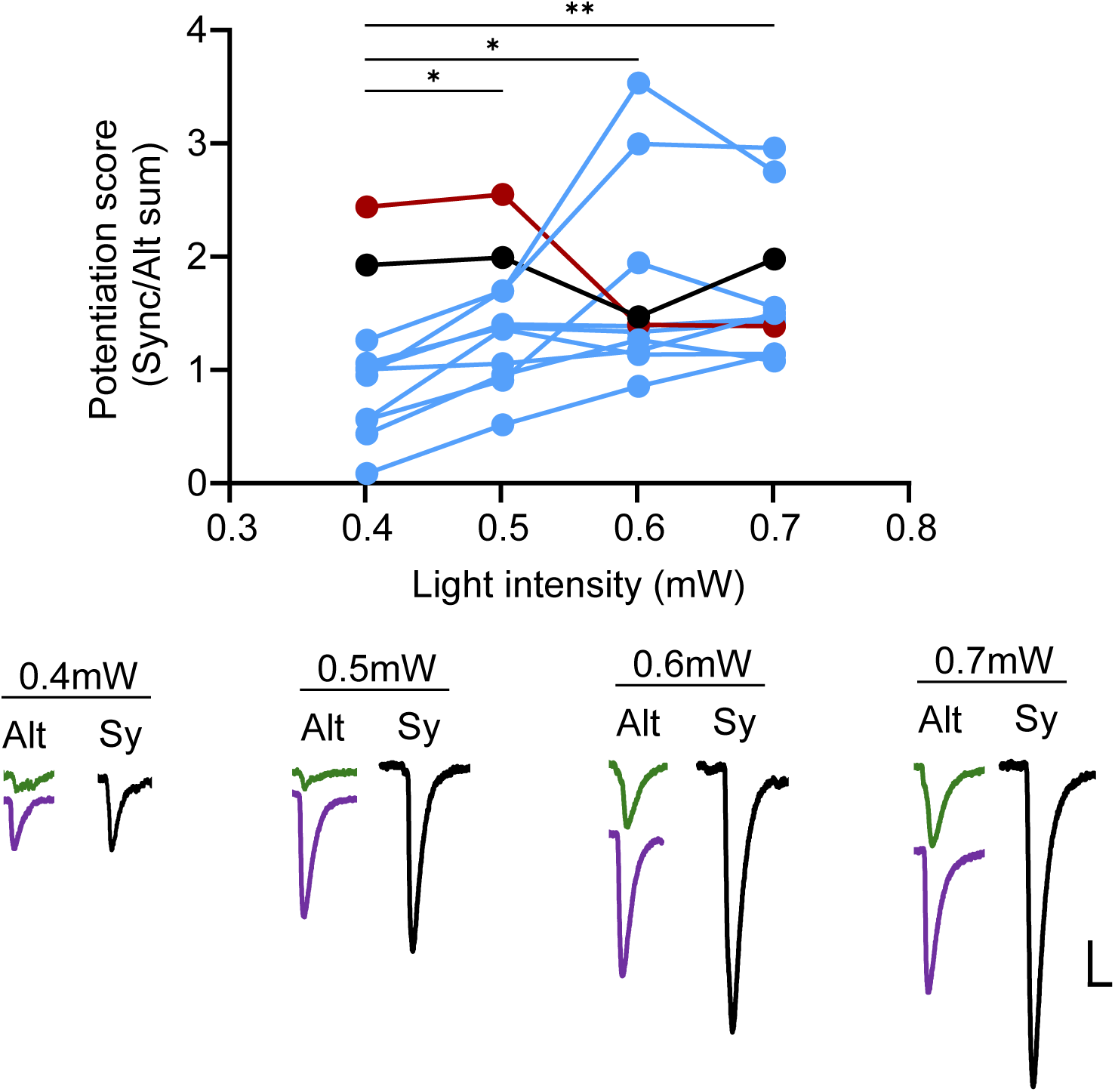
Heterosynaptic potentiation across multiple light intensities. Recording from dually innervated MSNs, sequential and synchronous stimulation assay reveals non-linear summation of responses at multiple light intensities (0.5-0.7mW). (n=11 cells/3 mice, p=0.0029, Friedman test with Dunn’s multiple comparisons test: 0.7 vs 0.4: **p=0.0031, 0.6 vs 0.4: *p=0.0300, 0.5 vs 0.4: *p=0.0494). Each line represents a cell. Black highlights a cell with no change across light intensities. Red highlights a cell where potentiation score decreased with increasing light intensity. Scale bars on representative traces = 20pA/20ms.

